# From atoms to cells: bridging the gap between potency, efficacy, and safety of small molecules directed at a membrane protein

**DOI:** 10.1101/2021.03.19.432117

**Authors:** Rodrigo Aguayo-Ortiz, Jeffery Creech, Eric N. Jiménez-Vázquez, Guadalupe Guerrero-Serna, Nulang Wang, Andre Monteiro da Rocha, Todd J. Herron, L. Michel Espinoza-Fonseca

## Abstract

Membrane proteins constitute a substantial fraction of the human proteome, thus representing a vast source of therapeutic drug targets. Indeed, newly devised technologies now allow targeting “undruggable” regions of membrane proteins to modulate protein function in the cell. Despite the advances in technology, the rapid translation of basic science discoveries into potential drug candidates targeting transmembrane protein domains remains challenging. We address this issue by harmonizing single molecule-based and ensemble-based atomistic simulations of ligand–membrane interactions with patient-derived induced pluripotent stem cell (iPSC)-based experiments to gain insights into drug delivery, cellular efficacy, and safety of molecules directed at membrane proteins. In this study, we interrogated the pharmacological activation of the cardiac Ca^2+^ pump (Sarcoplasmic reticulum Ca^2+^-ATPase, SERCA2a) in human iPSC-derived cardiac cells as a proof-of-concept model. The combined computational-experimental approach serves as a platform to explain the differences in the cell-based activity of candidates with similar functional profiles, thus streamlining the identification of drug-like candidates that directly target SERCA2a activation in human cardiac cells. Systematic cell-based studies further showed that a direct SERCA2a activator does not induce cardiotoxic pro-arrhythmogenic events in human cardiac cells, demonstrating that pharmacological stimulation of SERCA2a activity is a safe therapeutic approach targeting the heart. Overall, this novel platform encompasses organ-specific drug potency, efficacy, and safety, and opens new avenues to accelerate the bench-to-patient research aimed at designing effective therapies directed at membrane protein domains.

## Introduction

Membrane proteins are pivotal players in the cell, playing essential biological roles in a variety of functions vital to the survival of organisms, including transport ions and molecules, signal transduction across cells, serve as scaffolds to help bind the cell to a surface or substrate, and catalyze reactions in biological membranes ^1,2^. Membrane proteins constitute a significant fraction (about 20-30%) of the human proteome ^3^, and these proteins represent more than 60% of the current drug targets ^4^; enzymes, transporters, ion channels, and receptors are all common drug targets. Molecules directed at solvent-accessible pockets have been the primary strategy used to target membrane proteins ^5^; indeed, virtually all therapeutics targeting membrane proteins bind to solvated regions outside the lipid bilayer ^6^. Therefore, membrane protein-based drug discovery has rested on the primary assumptions that transmembrane domains are simply passive structural domains required to anchor membrane proteins to the lipid bilayer and that there are no specific sites and interactions within the transmembrane domains that can be effectively used for drug development. Recent advances in crystallography, spectroscopy and, computational biophysics now challenge this conventional view and show that transmembrane domains actively mediate functional protein-protein interactions and exert modulatory roles in membrane proteins ^7^. The paradigm shift from traditional structural biology to a dynamic view of membrane proteins has enabled the discovery of effector sites located in conventionally undruggable transmembrane regions to modulate protein function ^6,8,9^. Indeed, a prime example of a druggable transmembrane protein is the cardiac sarcoplasmic reticulum Ca^2+^-ATPase (SERCA2a). SERCA is an ATP-fueled pump that actively transports cytosolic Ca^2+^ into the sarcoplasmic reticulum during diastole in the heart, relaxing muscle cells and allowing the ventricles to fill with blood ^10^. SERCA2a regulation is critical for maintaining normal heart function, and its pathological dysregulation is a hallmark of heart failure. Stimulation of SERCA2a activity by allosteric modulators targeting the transmembrane domain results in improved cardiac function ^11-14^. Therefore, SERCA2a has become a promising pharmacological target for heart failure therapy ^15,16^, and also a new paradigm for membrane protein-based drug discovery ^14,17-19^.

Membrane permeation is considered the key factor limiting the utility of chemical probes and therapeutic agents ^20,21^, as the suboptimal ability of molecules to penetrate the cell and engage their membrane protein target results in poor efficacy. Nonetheless, it remains unclear whether membrane permeability profiles alone are sufficiently predictive to establish the relationships between target-based potency and cell efficacy of molecules targeting membrane protein domains. Another major limitation preventing the discovery of exogenous agents that can be used as selective probes that target transmembrane protein domains is the use of heterologous expression systems to evaluate cellular efficacy. This issue is of particular importance because small-molecule screening campaigns are often performed using cells that express the foreign target in quantities suitable for high-throughput screening ^22^ but do not necessarily capture organ-specific cellular target engagement in and potential off-target effects ^23^. Therefore, understanding the relationships between target-based potency and cell-based efficacy is of utmost importance to accelerate the discovery and development of biologically active molecules targeting transmembrane protein domains. To address this issue, here we used complementary single-molecule and ensemble–based atomistic simulations of lipid-ligand interaction, *in situ* enzymatic activity assays, and high-resolution optical mapping using human iPSC-derived cardiac cells to systematically interrogate pharmacological modulation of SERCA2a activation as a model. The result is a comprehensive platform that encompasses drug delivery, target engagement, cellular efficacy, and safety of molecules directed at membrane proteins.

## Results and Discussion

### Target-based potency of small molecules directed at cardiac SERCA2a

In this study, we used the only three known allosteric activators directed at the transmembrane domain of SERCA2a (**Figure 1**). CDN1163 was reported to activate the housekeeping SERCA2b isoform ^24^, whereas compounds CP-154526 and Ro 41-0960 have shown to activate cardiac SERCA2a isoform ^25^. Therefore, before testing our complementary computational and cell-based approach, we first evaluated the activity of these molecules directed at SERCA2a in cardiac sarcoplasmic reticulum (SR) microsomes isolated from hearts using an NADH-coupled ATPase assay. These ATPase activation assays confirmed that all three molecules stimulate SERCA2a activity with mean half-maximum effective concentration (EC_50_) values between 0.8-27 µM (**Table 1**). The target-based potency of these molecules *in situ* is either in the low micromolar (i.e., CDN1163 and Ro 41-0960) or high nanomolar range (i.e., CP-154526); therefore, these findings indicate that these molecules are relatively selective toward SERCA2a. The small molecules also increase ATPase turnover (V_max_) by 27-38% at a Ca^2+^ concentration of 10 µM (**Table 1**); these findings suggest that these SERCA activators stimulate maximal Ca^2+^ uptake rates in cardiac SR microsomal preparations. Together, these findings support previous studies and demonstrate that molecules CDN1163, CP-154526, and Ro 41-0960 are direct activators of the cardiac SERCA2a pump.

**Table 1.** Half-maximum effective concentration (EC_50_), change in ATPase turnover (V_max_), partition coefficient, and ligand-lipid free energy of binding of SERCA activators.

| Compound | EC<sub>50</sub> (μM)<sup>a</sup> | ΔV<sub>max</sub> (%)<sup>a</sup> | LogP<sub>BLM</sub> | ΔG<sub>bind</sub> neutral (kcal mol<sup>-1</sup>)<sup>b</sup> | ΔG<sub>bind</sub> ionized (kcal mol<sup>-1</sup>)<sup>b</sup> |
| --- | --- | --- | --- | --- | --- |
| CDN1163 | 1.6 ± 0.3 | 27 ± 3 | 2.3 | -3.4 | – |
| CP-154526 | 0.8 ± 0.4 | 31 ± 4 | 5.6 | -8.3 | -5.1 |
| Ro 41-0960 | 27 ± 1.3 | 38 ± 2 | -5.9 | -8.8 | -4.8 |

<sup>a</sup>Values were obtained from *in situ* SERCA2a ATPase activity experiments (*n*=3); values are reported as mean ± SEM.
<sup>b</sup>This value represents the energetic cost of burying the molecules within the bilayer was estimated using the PMF profile of the umbrella sampling calculations.

**Figure 1.**
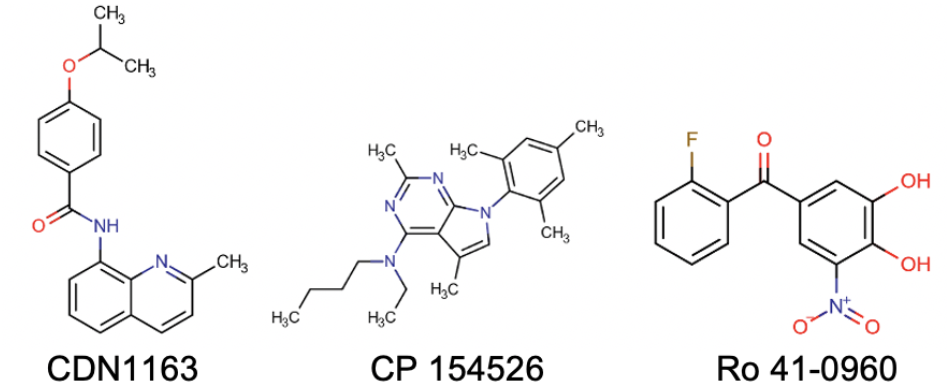
Structures of three activators known to target the transmembrane domain of the calcium pump.

### Bilayer crossing profiles of SERCA2a activators

Upon establishing the target-based potency of the three activators toward SERCA2a, we used atomistic simulations to evaluate the bilayer-crossing free energy profiles of these molecules. We predict that at physiological pH of 7.1-7.2,^26^ CDN1163 is electrically neutral (pKa= 4.0). Conversely, a pKa value of 6.4 calculated for CP-154526 (pK_a_=6.4) suggests that this molecule could be either neutral or ionized, whereas Ro 41-0960 (pK_a_=5.5) is predominantly charged at physiological pH. Based on these pK_a_ calculations, we computed the POPC bilayer-crossing free energy profiles of the CDN1163 and CP-154526 in their neutral state, and Ro 41-0960 in its ionized state. Additionally, we calculated the free-energy profiles for charged CP-154526 and neutral Ro 41-0960 to account for the uncertainty of the model (Root-mean square error of 1.1 pKa units) ^27^. The profiles in **Figure 2** highlight the range of energetic barriers possible with different SERCA2a activators and ionization states, with all the umbrella sampling windows provided in Figure S1 (Supporting Information). All electrically neutral molecules have a small membrane entry barrier from solution (**Fig. 2**), but the energy barrier is not considerably affected by ionization, as observed in the bilayer-crossing free energy profiles of electrically charged CP-154526 and Ro 41-0960 (**Fig. 2**). After crossing this energetic barrier, the compounds reach their preferred penetration depth, either close to the lipid headgroup or the membrane midplane (**Fig. 2**). Except for the ionized states of CP-154526 and Ro 41-0960, there is a peak for the free energy profile at the center of the bilayer representing the energetic cost of burying the molecules within the bilayer center. Despite these differences, all molecules interact favorably with the lipid bilayer regardless of their ionization state, with free energies of binding (ΔG_bind_) between −3.4 and −8.8 kcal mol^-1^ (**Table 1**). More importantly, we found that Ro 41-0960 has a high affinity for the lipid bilayer, in contrast to the predicted negative logP_BLM_ calculated for this molecule (**Table 1**).

**Figure 2.**
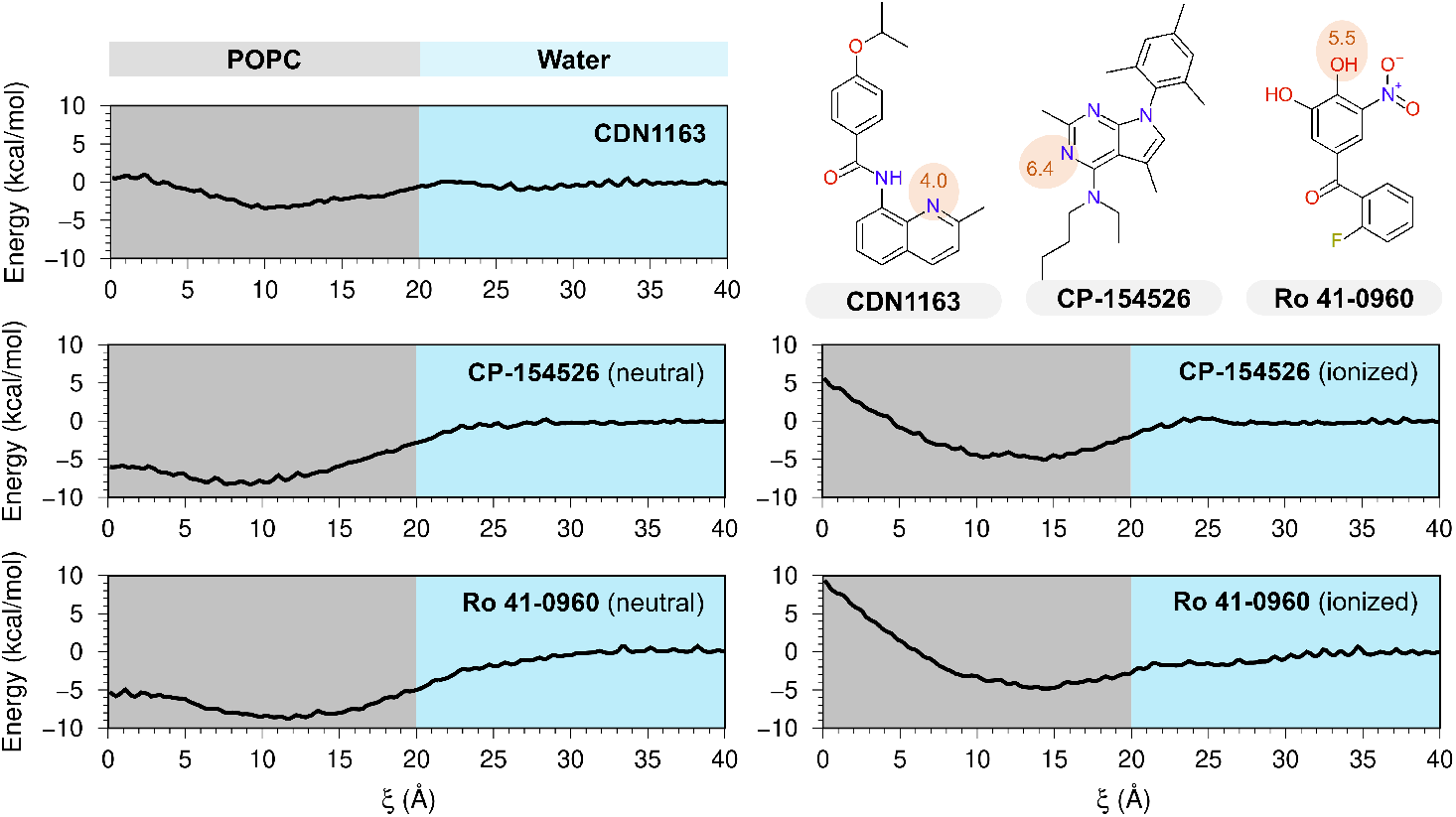
Bilayer-crossing free energy profiles of SERCA2a activators. Profiles are represented by the potential of the mean force (PMF) of SERCA2a activators CDN1163, CP-153526, and Ro 41-0960 along the normal z-axis of a POPC membrane; the distance along the z-axis was used here as a reaction coordinate (ξ). The chemical structures in the upper right indicate the location of titratable groups and their corresponding pKa values (red circles); based on these pKa values, Bilayer-crossing free energy profiles were calculated for either neutral or ionized states of the small molecules.

### Single-molecule simulations of ligand-membrane interactions

We performed unrestrained molecular dynamics (MD) simulations of the activators embedded in a POPC bilayer as a complementary approach to evaluate the stability of the ligand-membrane interactions. **Figure 3** shows the time-dependent and probability distribution of the vertical position of the molecules in the lipid bilayer, where [Z] = 0 Å denotes the membrane center of mass. In agreement with the bilayer-crossing free energy profiles, unrestrained MD simulations show that the molecules remain bound to the lipid bilayer and localize at the interface between the polar and nonpolar regions of the membrane, e.g., near or below the headgroups of POPC (**Figure 3**). The vertical position of the molecules is determined by the charge of the compounds: electrically neural molecules penetrate deeper into the membrane at distances between 9-12 Å from the membrane center of mass, whereas charged molecules localize farther from the center of the lipid bilayer (**Figure 3**). For example, CDN1163 binds about 9 Å away from the center of the bilayer, while Ro 41-0960 maintains an average distance of 14 Å from the center of mass of the POPC bilayer and can further localize into the headgroup-water interface (e.g., at *t*=25-35 ns, **Figure 3**). The unrestrained MD simulations correlate well with PMF calculations and show that both electrically neutral and charged states of Ro 41-0960 remain bound to the lipid bilayer despite carrying a charged nitro group. The qualitative and quantitative agreement between biased and unbiased atomistic simulations imply that these activators interact favorably with a lipid bilayer. The affinity of these molecules for the lipid bilayer supports the concept that these effectors activate SERCA2a by interacting with the transmembrane domain of the protein.

**Fig. 3.**
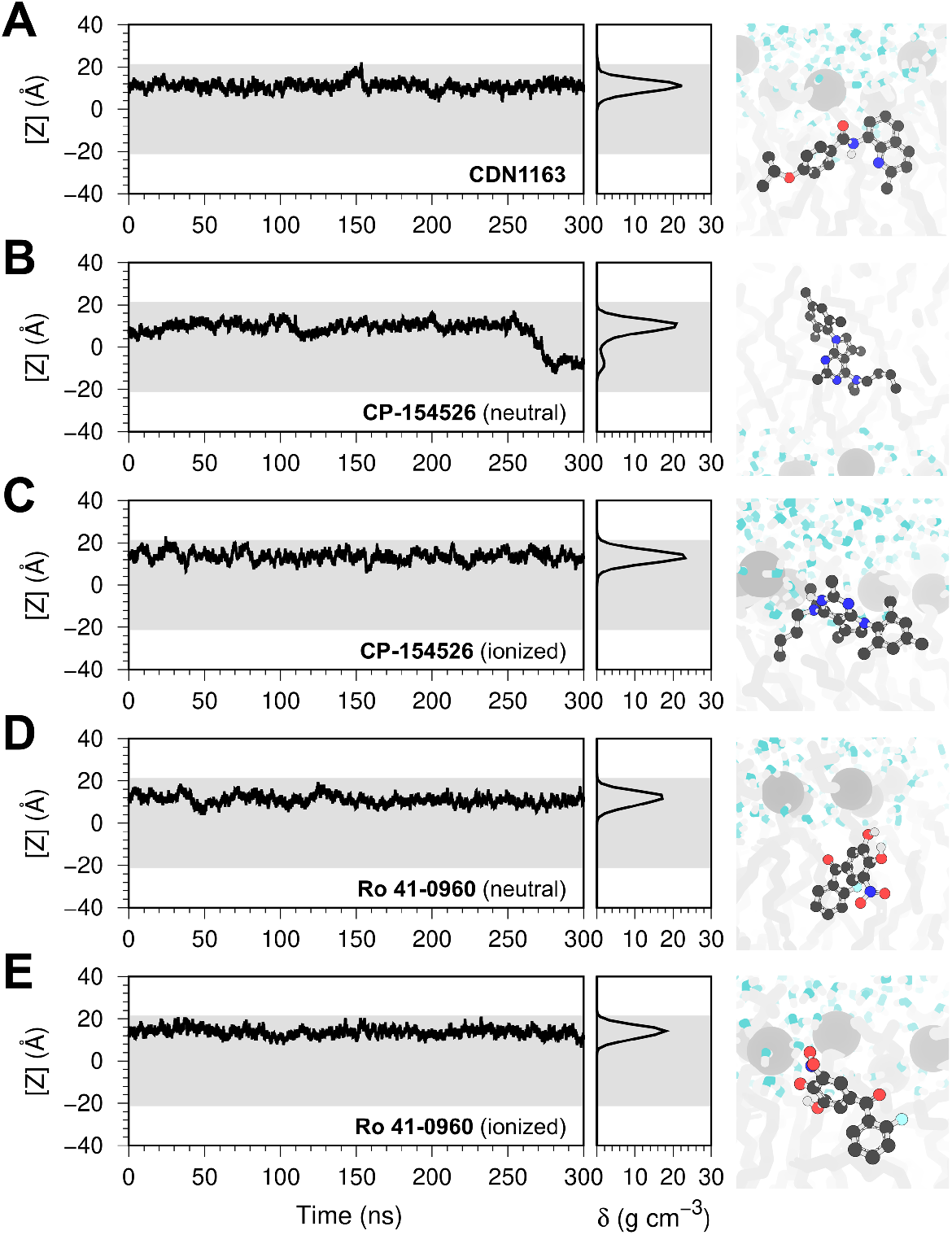
Single-molecule atomistic simulations of ligand-membrane interactions. The plots represent the position of a single molecule of (A) CDN1163, (B, C) CP-154526, and (D, E) Ro 41-0960 bound to a solvated POPC membrane. We show the time-dependent position of the molecule along the z-axis of the lipid-water system (left), the relative density (δ) profile of the molecules in the lipid bilayer (center), and the orientation of the molecules in the lipid bilayer extracted at the end of the trajectories (right). In all plots, the shaded area represents the location of the lipid bilayer. The small molecules are shown as ball and stick representation; lipids and water molecules are shown in gray and blue, respectively.

In addition to the localization in the lipid bilayer, another important variable to consider is the translocation of the small molecules across the membrane. This is especially important for targets localized in intracellular compartments, including the sarcoplasmic reticulum where SERCA2a is found. Lipid-bilayer crossing free-energy profiles show that electrically neutral CP-154526, Ro 41-0960, and to a certain extent CDN1163 can translocate freely across the lipid bilayer. Analysis of the unbiased MD trajectories showed that electrically neutral CP-154526 undergoes a single spontaneous exchange event between membrane leaflets (i.e., at *t*=275 ns, **Fig. 3**). Although the PMF profiles of electrically neutral Ro 41-0960 and CDN1163 indicate that these molecules can cross the membrane, we did not observe leaflet exchange of this molecule in the timescales studied here. Based on the lipid-ligand and bilayer-crossing energy profiles, we can infer that CP-154526 will elicit the highest effect on stimulation of Ca^2+^ transport, CDN1163 will produce a moderate pharmacological response, and Ro 41-096 will produce a limited effect on Ca^2+^ dynamics in cardiac cells.

### Cell-based efficacy of SERCA2a activators in human iPSC-derived cardiac cells

We determined the concentration-response effects on Ca^2+^ flux dynamics and beating rate in spontaneously beating human iPSC-derived cardiomyocytes using high-resolution optical mapping. Human iPSC-derived cardiomyocytes used here are ideally suited for these studies because they express all major components of the Ca^2+^ transport machinery (**Fig S2-A**, Supplementary Information) and respond to β-adrenergic-induced phosphorylation phospholamban, the main regulator of SERCA2a activity in the heart (**Fig. S2-B**, Supplementary Information). The efficacy of SERCA2a activation *in vitro* is determined by the rise and decay in cytosolic Ca^2+^ (the Ca^2+^ transient amplitude), the Ca^2+^ transient duration at 50% recovery (CaTD_50_), and by changes in chronotropy. Pharmacological activation of SERCA2a is expected to stimulate Ca^2+^ cycling in the cardiomyocyte, as reflected by the ability of small molecules to increase the Ca^2+^ transient amplitude and decrease the CaTD_50_ ^14,28^. Treatment of human iPSC-derived cardiomyocytes with CP-154526 increases Ca^2+^ transient amplitude in a concentration-response manner; however, the effect here follows a bell-shaped behavior, reaching a maximum increase in the Ca^2+^ transient amplitude (23%) at compound concentrations of 1-5 µM (**Fig 4A**). CP-154526 produces contrasting effects on CaTD_50_ at various concentrations; for instance, CP-154526 decreases CaTD_50_ by 9% at a compound concentration of 0.1 µM, while it increases CaTD_50_ by 6% at a concentration of 5 µM (**Fig 4A**). More importantly, CP-154526 has a substantial effect on chronotropy, increasing the beat rate of cardiac cells by 16-33% at compound concentrations 0.1, 50, 10, and 50 µM (**Fig. 4A**). CP-154526 generally slows down intracellular Ca^2+^ dynamics and also has a positive chronotropic effect. These findings are unexpected because the effects produced by CP-154526 are opposite to those exerted by direct stimulation of SERCA2a activity in human cardiac cells ^14,28^. Conversely, CDN1163 displays a concentration-response profile that is consistent with the stimulation of SERCA2a activity in the cardiac cells ^14,28^: a concentration-dependent increase in the Ca^2+^ transient amplitude with a maximum effect at a concentration of 0.5 µM, and a decrease in CaTD_50_ that is consistently induced by the compound at concentrations 0.5 through 50 µM (**Fig 4B**). Unlike CP-154526, we found that CDN1163 has negligible chronotropic effects on human iPSC-derived cardiomyocytes at all concentrations tested in this study (**Fig. 4B**). In agreement with our original hypothesis, Ro 41-0960 has a marginal effect on the Ca^2+^ transient amplitude at concentrations equal to or less than 10 µM (**Fig. 4C**). Nevertheless, Ro 41-0960 produces a moderate effect on the CaTD_50_ (**Fig. 4C**). Interestingly, Ro 41-0960 exerts contrasting effects on the Ca^2+^ transient amplitude and CaTD_50_ at low and high concentrations: at concentrations lower than 10 µM, Ro 41-0960 increases Ca^2+^ transient amplitude and CaTD_50_, whereas the opposite effects are induced at compound concentrations of 50 and 100 µM (**Fig. 4C**). Ro 41-0960 does not generally affect the cardiomyocyte beat rate, although a modest negative chronotropic effect is observed at compound concentrations of 0.5 and 10 µM (**Fig. 4C**).

**Fig. 4.**
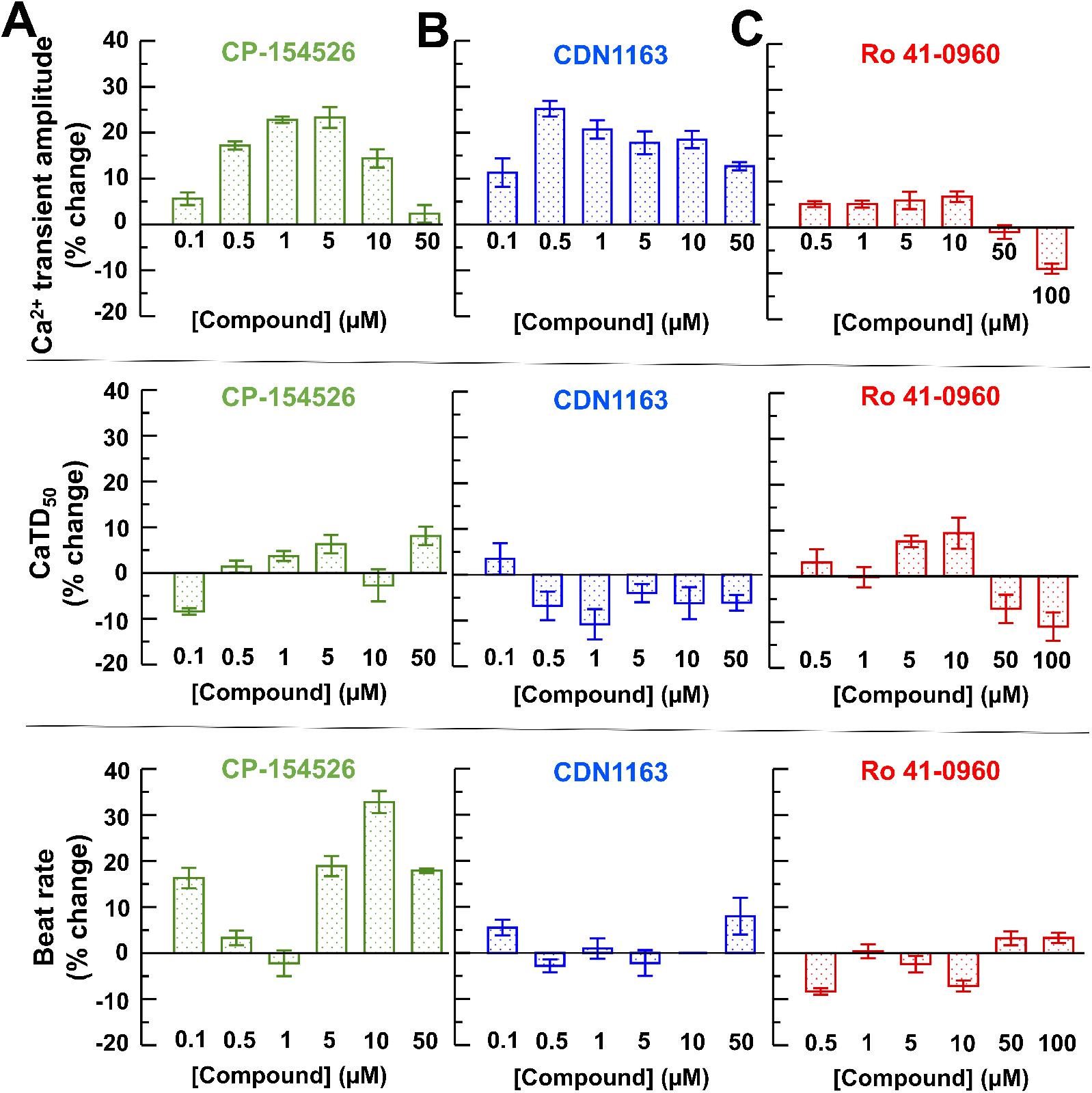
Functional characterization of SERCA2a activators in human iPSC-derived cardiomyocytes. Concentration-response curves were obtained for the effects of (A) CP-154526 (B) CDN1163, and (C) Ro 41-0960 on the percentage of change in the Ca^2+^ transient amplitude, Ca^2+^ transient duration at 50% recovery (CaTD_50_) and beat rate in normal human iPSC-derived cardiomyocytes. Results are expressed as percent change from baseline values in Hank’s balanced salt solution and reported as mean ± SEM (*n*=6).

### Ensemble-based atomistic simulations of ligand-membrane provide unique insights into ligand-membrane interactions

In agreement with our initial simulation-based hypothesis, human iPSC-derived cardiomyocyte assays indicate that CDN1163 engages SERCA2a in the cardiomyocyte and that the bilayer-crossing energy profile of Ro 41-0960 is suboptimal for delivery into the cell. However, the cell-based activity of CP-154526 is in contrast with its nanomolar target-based potency and optimal lipid-ligand interaction profiles for intracellular delivery and recruitment into the SR membrane. To explain this contradictory evidence, we complemented our biased and unbiased single-molecule approaches with ensemble-based atomistic simulations ^29^. These ensemble-based simulations provide an atomistic representation of ligand-ligand/ligand-lipid interactions and ligand translocation across the membrane in real-time. We found that ligand-lipid interactions are reproducible across biased and unbiased MD approaches. Specifically, the MD simulations of the systems containing multiple SERCA2a activators capture the spontaneous insertion of small molecules into the membrane (**Fig. 5**). The insertion depth found in these simulations is in good quantitative agreement with the biased and unbiased MD simulations. For instance, the maximum relative density of CDN1163 in the lipid bilayer occurs at 11-12 Å from the center of the bilayer, whereas electrically charged Ro 41-0960 resides primarily at ∼14 Å from the center of the bilayer (**Fig 6**). The relative density profiles of the CP-154526, especially in its electrically charged state, are also in good qualitative and quantitative agreement with unbiased single-molecule MD simulations (**Fig. 3**). The reproducibility of ligand-membrane interactions across different MD approaches validates the ensemble-based simulations as an approach to gain further insights into the behavior of small molecules in a lipid-water environment.

**Fig. 5.**
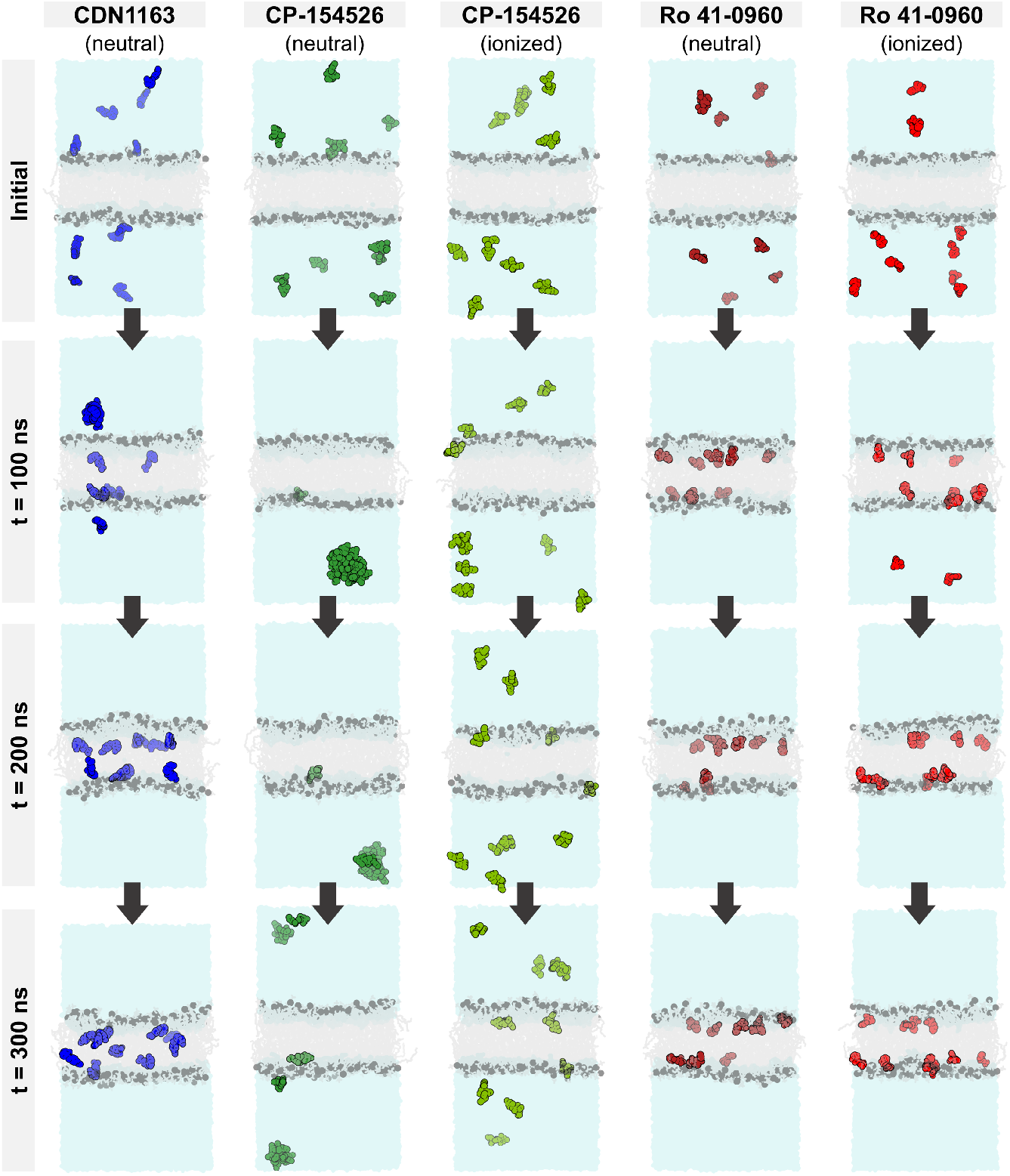
Time-resolved behavior of SERCA2a activators in a lipid-water environment. Each snapshot represents a configuration extracted from the ensemble-based atomistic simulations of a system containing a solvated membrane and ten copies of each SERCA2a activator. The simulations capture spontaneous insertion of small molecules into the membrane as well as ligand-ligand interactions during the 300 ns of simulation time. CDN1163 (blue), CP-154526 (green), and Ro 41-0960 (red) are shown as spheres; lipids and water molecules are shown in gray and blue, respectively.

**Fig. 6.**
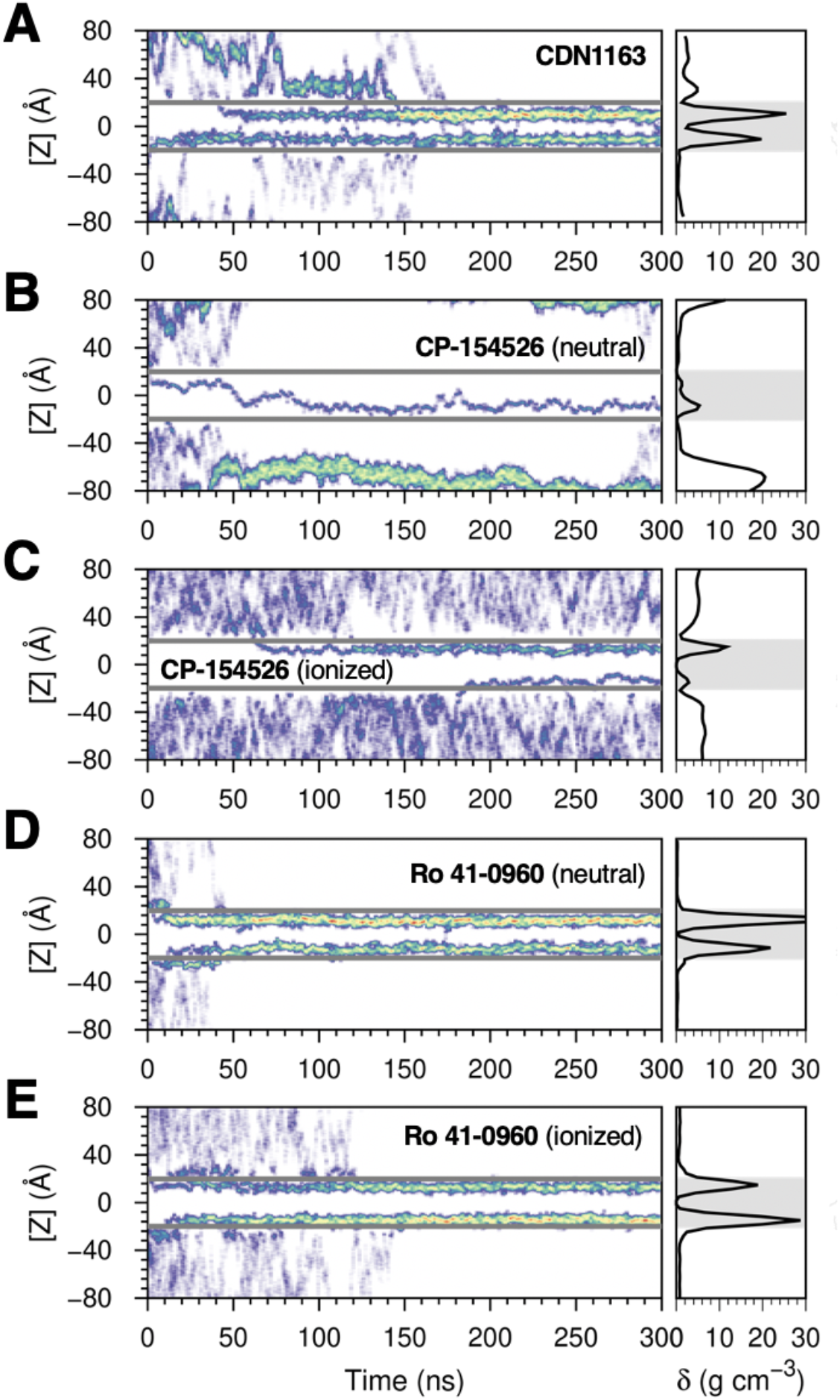
Relative density profiles of SERCA2a activators in a lipid-water system obtained from the ensemble-based MD simulations. The figure shows the position of the molecules relative to the z-axis of the system (left) and the relative density (δ) profile (right) calculated for (A) CDN1163, (B, C) CP-154526, and (D, E) Ro 41-0960. The gray horizontal lines (left) and the shaded area (right) represent the boundaries of the lipid bilayer.

Further inspection of the relative density profiles obtained from the ensemble-based simulations of CDN1163 showed that the 10 molecules of CDN1163 simulated in the lipid-water system bind to the lipid bilayer at t=170 ns and remain bound to the membrane for the remainder of the simulation time (**Fig. 6A**). In contrast with the favorable ligand-lipid interaction single-molecule profiles of CP-154526 (**Figs. 1 and 2**), ensemble-based simulations showed that this molecule has a strong preference for the aqueous phase of the system (**Fig. 6B, C**). Surprisingly, the preference for the aqueous phase is more pronounced in the electrically neutral state of the molecule, where only 10-20 % of the CP-154526 molecules transfer to the lipid bilayer (**Fig. 6B**). More importantly, we found that electrically neutral CP-154526 aggregates in the aqueous phase (**Fig 6B**), although ionization partially mitigates this effect (**Fig. 6C**). Finally, MD simulations of Ro 41-0960 showed that all 10 molecules bind to the lipid bilayer at t=40 ns for the electrically neutral molecule and t=140 ns for the electrically charged form of the ligand; as in CDN1193, Ro 41-0960 remains bound to the lipid bilayer in both trajectories. We found that this behavior is consistent across ionization states, thus indicating that changes in ionization state do not affect the ability of Ro 41-0960 to associate with the lipid bilayer. In addition to the differences in ligand-ligand and ligand-lipid interactions observed in the ensemble-based simulations, a notable discovery made here is the clear differences in ligand translocation profiles across the lipid bilayer that are not entirely captured by single-molecule simulations. Specifically, the density profile of CDN1163 shows an increase in the relative density of this molecule around the center of the lipid bilayer ([*Z*] = 0 Å, **Fig 6A**), suggesting that this molecule penetrates across the lipid bilayer in the time scales used in this study. Similarly, electrically neutral CP-154526 penetrates across the membrane; penetration occurs only once in the trajectory, which is explained by the preference of this molecule to partition into the aqueous phase. Conversely, the relative density of the small molecules around the center of the lipid bilayer is nearly 0 g cm^-3^ in the trajectories of electrically charged CP-154526 as well as Ro 41-0960 (charged and neutral), thus indicating that these molecules do not penetrate across the lipid bilayer in the MD trajectories (**Fig. 6D, E**).

### Atomistic simulations and cell-based experiments bridge the gap between potency and efficacy of activators targeting SERCA2a

An important question arises from the studies presented here: How do simulations and experiments integrate to bridge the gap between target-based potency and efficacy of bioactive molecules directed at the transmembrane protein? To answer this question, it is instructive to pick two similar examples: CDN1163 and CP-154526 activate SERCA2a *in situ* with similar potency, yet these molecules produce different effects in human iPSC-derived cardiac cells, with only CDN1163 producing a pharmacological response expected as a result of direct SERCA2a activation. Here, the ensemble-based MD simulations provide a mechanistic explanation for these differences: while both molecules can penetrate across the membrane, CDN1163 is expected to engage SERCA2a more efficiently as it is predominantly bound to the lipid bilayer. Conversely, the preference of CP-154526 for the aqueous phase lowers its delivery of the SR membrane, thus reducing the engagement of SERCA2a in the cardiomyocyte. Therefore, the differences in cell-based efficacy may be attributed to a combination of suboptimal ligand delivery to the lipid bilayer, as detected by our MD simulations, as well as off-target effects, as suggested by our cell-based screening assays. It is conceivable that the pharmacological effects of CP-154526 on Ca^2+^ dynamics and chronotropy observed in human iPSC-derived cardiomyocytes results from ligand-protein interactions with soluble proteins outside the sarcoplasmic reticulum ^30,31^. Moreover, the self-aggregation of the CP154526 observed in the ensemble-based simulations (**Fig. 5**) also explains the bell-shaped concentration-response curve obtained in the iPSC-based experiments (**Fig. 4A**) ^32^. Based on this evidence, we can conclude that CDN1163, and not CP-154526, activates SERCA2a activity in human iPSC-derived cardiac cells. This complementary approach also explains the effects of Ro 41-0960 on the cardiomyocyte: this molecule activates SERCA2a in isolated cardiac SR microsomes, in agreement with simulations showing that Ro 41-0960 binds efficiently to a lipid bilayer. However, high-resolution optical mapping experiments showed that this molecule induces a marginal effect on the intracellular Ca^2+^ dynamics of human iPSC-derived cardiac cells, in agreement with atomistic simulations showing that Ro 41-0960 permeates poorly across the cell membrane. Nevertheless, we speculate that the bilayer partitioning of Ro 41-0960 facilitates its non-specific interaction with ion transporting channels found in the plasma membrane ^33^. This off-target effect is supported by high-resolution optical mapping experiments showing that at concentrations of 50-100 µM, Ro 41-0960 produces changes in the Ca^2+^ dynamics that are similar to those induced by Ca^2+^ channel blockers ^34^.

It is important to note that while CP-154526 and Ro 41-0960 do not produce the expected pharmacological effects on human iPSC-derived cardiac cells, the high-throughput screening campaigns that led to the discovery of these molecules are not ‘wrong’ ^25^, because these molecules are actual activators of SERCA2a *in situ* (**Table 1**). Instead, these molecules can be integrated into future structure-activity campaigns aimed at (i) elucidating transmembrane effector sites that can be targeted for therapeutic modulation (ii) designing new chemical probes with optimal potency and cell-based efficacy, and (iii) dissecting the molecular mechanisms underlying small-molecule SERCA activation. Achieving these goals will ultimately enable advance therapeutic interventions that directly target the molecular basis of the disease in the failing heart ^35^.

### Pharmacological activation of SERCA2a does not induce pro-arrhythmogenic events in cardiac cells

Finally, we addressed drug-induced cardiac toxicity, an issue of significant concern in drug discovery trials, particularly when a promising new lead has a desired pharmacological effect that occurs at the expense of long-term pro-arrhythmia cardiotoxicity ^36,37^. Hence, an important question we ask here is whether pharmacological activation of SERCA2a may induce Ca^2+^-dependent changes in excitability that lead to arrhythmogenic currents originating from the overload of SR with Ca^2+^ in response to sustained stimulation of the pump ^38^. To rule out this effect, we performed drug-induced cardiotoxicity screening using high-resolution optical mapping with a voltage-sensitive dye to test the effects of CDN1163 on conduction velocity (CV, **Fig 7A**) and action potential duration at 90% repolarization (APD_90_) in spontaneously beating human iPSC-derived cardiac cells (**Fig 7B**). We found that neither CV nor APD_90_ was significantly affected by the treatment of cardiomyocytes with the SERCA activator CDN1163. Specifically, we found that the CV was 17.5 ± 5 cm/s for untreated cells compared to 19.3 ± 4 cm/s, 25.6 ± 8 cm/s, and 16 ± 3 cm/s for treatment of cells with 0.5, 5 and 50 μM concentrations of CDN1163, respectively (**Fig. 7C**). Interestingly, we found that the CV is not compromised in electrically coupled iPSC-CMs monolayer syncytia even at very high concentrations of the compound. Furthermore, the APD_90_ values were similar in all experimental conditions: 324.2 ± 13 ms for untreated cells; and 326.2 ± 31 ms, 342.9 ± 31 ms, and 344.2 ± 24 ms for monolayer syncytia treated with CDN1163 at concentrations of 0.5, 5, and 50 μM, respectively (**Fig. 7D**). Altogether, the data give us an idea that the compound CDN1163 is not leading to significant electrical alterations in iPSC-derived cardiomyocytes. These findings are important because changes in CV and/or APD_90_ are associated with pro-arrhythmia effects, such as re-entrant excitation or early/delayed afterdepolarizations ^37,39,40^. Therefore, our data indicate that pharmacological stimulation of SERCA2a activity using a relatively selective activator does not induce pro-arrhythmogenic effects in human iPSC-derived cardiac cells. These studies illustrate the importance and feasibility of integrating safety studies earlier into drug discovery pipelines aimed at validating the use of SERCA2a and membrane proteins in general as effective therapeutic targets.

**Figure 7.**
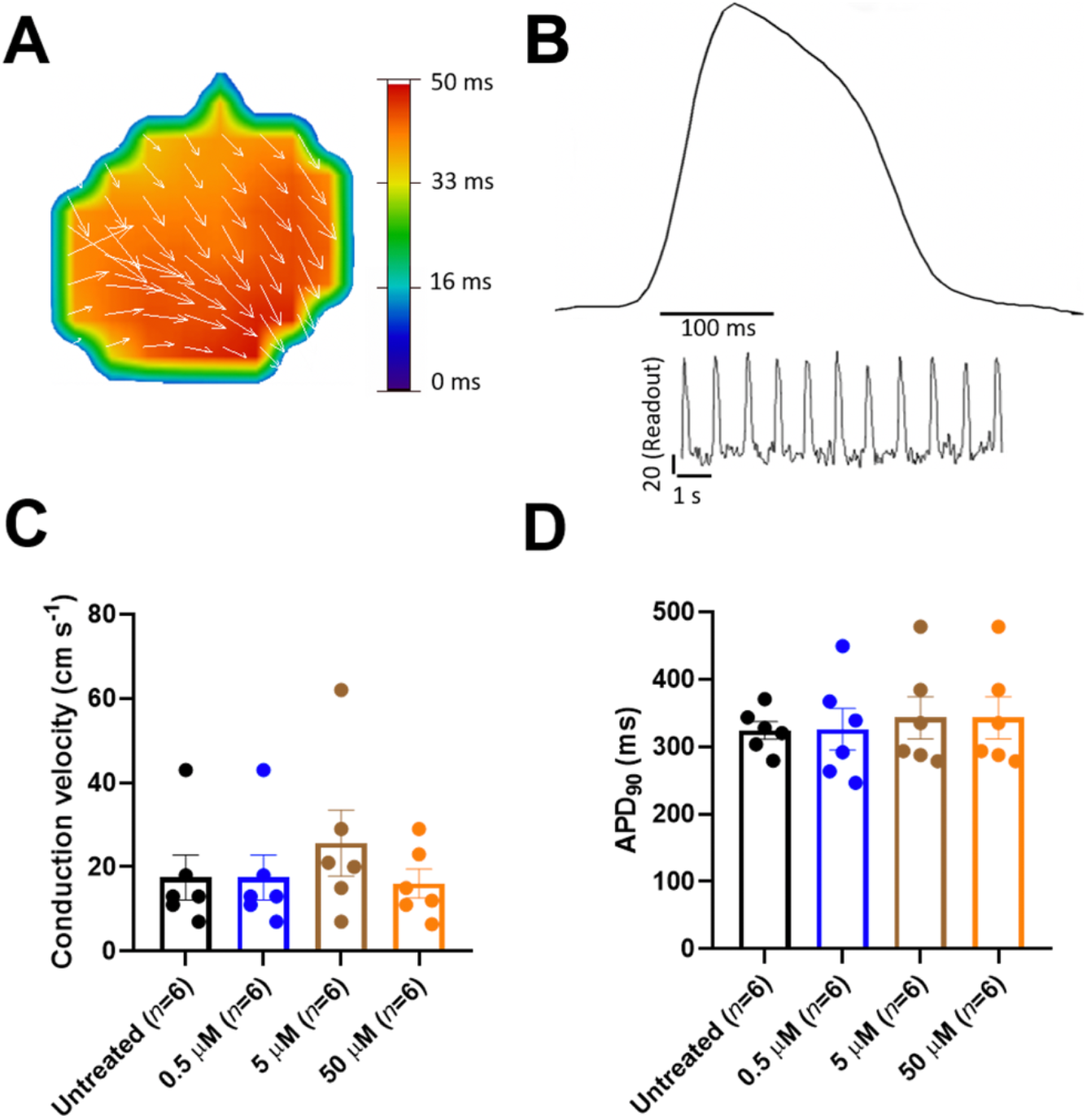
Conduction velocity and APD_90_ after treatment of human iPSC-derived cardiomyocytes with the SERCA activator CDN1163. (A) Representative impulse propagation during the spontaneous activity of untreated monolayer syncytium formed of human iPSC-derived cardiac cells. Impulse propagation is represented by the activation maps of action potential propagation at different activation times, with the arrows showing the direction of the propagation wave across the monolayer syncytium. (B) Representative trace of the spontaneous action potential (AP) obtained by optical mapping of a syncytium formed by human iPSC-derived cardiomyocytes. The inset represents single-pixel signals of spontaneous optical AP recordings. (C) Conduction velocity (CV) calculated from each activation map in the presence and absence (untreated control) of CDN1163. We found no significant changes in CV between the control and treatment groups. (D) Quantification of APD_90_ from optical mapping readings shown in panel ‘B’. We found that treatment of human iPSC-derived cardiomyocytes with CDN1163 does not have significant effects on the APD90 compared to the untreated control. Data are reported as mean ± SEM (*n*=6). Groups were compared using a one-way analysis of variance (ANOVA) with Dunnett test; statistical significance was set at *p*<0.05.

## Conclusion

In summary, we studied small-molecule activation of SERCA2a in human cardiomyocytes to explain the relationships between target-based potency and cell-based efficacy of bioactive molecules targeting a transmembrane protein domain. This approach, which integrates atomistic simulations and high-resolution experiments using human iPSCs, provides a vivid picture of cellular drug delivery and target engagement of molecules directed at membrane proteins. The significance of this approach is illustrated by its potential to distinguish and explain selective *vs* off-target effects early in the drug discovery process as well as its broad applicability to a variety of organ-and patient-specific human cell lines ^41,42^ and complementary experimental and computational tools ^43-45^. The result is a powerful platform to accelerate the discovery and design of bioactive molecules with well-defined potency, selectivity, and cell permeability that can be applied for the systematic interrogation and pharmacological modulation of membrane protein function. The flexibility of this platform to use iPSC allows it to expand into the areas of precision medicine, as it allows the use of patient-derived cells and organoids for therapeutic discovery ^46^. This platform can also be extended to comprehensive ‘clinical trials in a dish’ to test therapeutic agents for efficacy and safety in the laboratory with human tissue ^47^.

## Materials and Methods

### Compounds setup and LogP prediction

The 3-D structures of SERCA2a activators CDN1163 (CID: 16016585), CP-154526 (CID: 5311055), and Ro 41-0960 (CID: 3495594) were retrieved from the PubChem database (https://pubchem.ncbi.nlm.nih.gov) ^48^. Geometry optimization was performed using the MMFF94s force field implemented in the *obminimize* module of OpenBabel ^49^. We employed the chemicalize tool developed by ChemAxon ^50,51^ to predict the p*K*_a_ of the three molecules. The passive translocation of each of the compounds across a lipid bilayer of 1,2-dioleoyl-*sn*-glycero-3-phosphocholine at 300 K and pH 7.1 was carried out with the PerMM server ^52^ using “Drag” method to calculate the pathway. We assigned the pKa values and the charge at neutral pH in the PDB header of compounds CP-154526 (p*K*_a_ = 6.37, basic N atom) and Ro 41-0960 (p*K*_a_ = 5.54, acidic oxygen atom). From this calculation, we retrieved the permeability coefficient of molecules through the black lipid membrane (logP_BLM_).

### Bilayer crossing profiles of SERCA2a activators

Umbrella sampling simulations were carried out across a 1-palmitoyl-2-oleoyl-*sn*-glycero-3-phosphocholine (POPC) membrane for each of the compounds with the AMBER99SB-ILDN force field^53^ implemented in GROMACS 5.1.4 ^54^. Ligand topologies and parameters were generated with the ACPYPE interface^55^ using AM1-BCC method to compute partial charges. POPC lipid parameters were taken from the Slipids (Stockholm Lipids) force field ^56^. For the system setup, we set the molecules approximately 4.0 nm away from the center of mass of a pre-equilibrated lipid bilayer comprised of 100 POPC molecules. The system was solvated using the TIP3P water model and neutralized with sodium and chloride ions. Energy minimization and equilibration using NVT and NPT ensembles (1 ns e/a) were carried out restraining the initial position of the molecule in the aqueous solution. The equilibrated systems were further used to pull the compound into the center of the POPC lipid bilayer during 500 ps with a pulling rate of 0.01 nm ps^-1^ and a harmonic force constant of 500 kJ mol^-1^ nm^-2^. The pulling simulation was performed at 310 K and 1.0 bar using the Nosé-Hoover thermostat ^57^ and the semi-isotropic Parrinello–Rahman barostat ^58^. Forty-three to forty-six configurations were selected along the z-axis reaction coordinate (ξ) for the umbrella sampling simulations. Each configuration was energy minimized and submitted to a restrained NPT equilibration before 10-ns of NPT production run applying a harmonic force constant of 1000 kJ mol^-1^ nm^-2^ to the compound. The position and force data of all the configurations were evaluated with the Weighted Histogram Analysis Method (WHAM) ^59^ to generate the potential of mean force (PMF) profile. The snapshot with the lowest energy in the PMF profile of each compound was submitted to 300 ns non-restrained NPT production runs at 310 K and 1.0 bar using the previously described thermostat and barostat algorithms. The density profiles were computed with the *density* built-in tool of GROMACS 5.1.4, while the center of mass position of the molecule in the z-axis of the membrane was calculated with MDAnalysis python library ^60^.

### Ensemble-based MD simulations of unbound molecules in a lipid-water environment

A total of 10 molecules of each compound, to a final concentration of ∼17 µM, were randomly placed in the water phase of a system containing a POPC lipid bilayer consisting of 250 lipid molecules and about 34,300 water molecules. The system was energy minimized and equilibrated during 1 ns using NVT and NPT ensembles. The equilibrated system was subsequently submitted to 300 ns NPT production run using the previously described parameters. Graphs were made using Gnuplot 5.0 61; visualization was performed using PyMOL version 1.7 ^62^.

### Obtention of cardiac muscle sarcoplasmic reticulum vesicles containing SERCA2a

Pig hearts were obtained post-mortem from a near abattoir and placed in ice-cold 10 mM NaHCO_3_. Ventricular tissue was isolated, minced, and homogenized with a Waring immersion blender in 5 volumes of 10 mM NaHCO_3_ at 4°C. All procedures were done at 4°C and all the buffers contained protease inhibitors (Sigma, St. Louis, MO, USA). The homogenate was centrifuged at 6,500 *g* for 20 min. The supernatant was filtered through 6 layers of gauze and centrifuged at 14,000 *g* for 30 min. The supernatant was filtered and centrifuged at 47,000 *g* for 60 min. The pellet was resuspended in a buffer containing 0.6 M KCl and 20 mM Tris (pH = 6.8) using a Teflon Potter-Elvehjem homogenizer. The suspension was then centrifuged at 120,000 *g* for 60 min. The pellet obtained was resuspended and homogenized in a solution containing 0.3 M sucrose and 5 mM HEPES (pH 7.4). The protein concentration of the microsomal suspension was determined by using Coomassie Plus assay kit (Thermo Fisher Scientific, Waltham, MA, USA) in a Synergy H1 multi-mode plate reader (BioTek, Winooski, VT, USA).

### Measurement of SERCA2a ATPase activity

Concentration-dependent SERCA2a activation assays of CD1163, CP-154526, and Ro 41-0960 (Sigma-Aldrich, St. Louis MO, USA) were performed using microsome preparations obtained from fresh pig hearts. An enzyme-coupled, NADH-linked ATPase assay was used to measure SERCA2a ATPase activity in 96-well microplates. Each well contained assay mix (50 mM MOPS, pH 7.0), 100 mM KCl, 5 mM MgCl_2_, 1 mM EGTA, 0.2 mM NADH, 1 mM phosphoenolpyruvate, 10 IU/mL of pyruvate kinase, 10 IU/mL of lactate dehydrogenase, and 1 µM of the calcium ionophore A23187 (Sigma-Aldrich, St. Louis, MO, USA), and added CaCl_2_ to set the free Ca^2+^ concentration to 10 µM. 4 µg/mL of microsomal preparation, CaCl_2_, compound, and assay mix were incubated for 20 minutes. We tested activity at 11 different concentrations of the compound in the range of 0.05–50 µM (*n*=3 per concentration). The assay was started upon the addition of ATP, to a final concentration of 5 mM, and absorbance read at 340 nm in a Synergy H1 multi-mode plate reader (BioTek, Winooski, VT, USA).

### Culture of human iPSC-derived cardiomyocytes

We used cryopreserved human iPSC-derived cardiomyocytes (iCell^®^Cardiomyocytes^2^, FUJIFILM Cellular Dynamics, Madison, WI, USA). The cells were thawed and plated on custom-made 96-well or 6 well plates with inserts of polydimethylsiloxane (PDMS). PDMS vulcanized silicone transparent sheeting was obtained from SMI (Specialty Manufacturing, Inc, Saginaw, MI, USA) with 40D (D, Durometer or ≈1000 kPa) hardness and coverslips were cut out to fit in each well of 96 well or 6 well plates. PDMS coverslips were manually added to each well of 96/6 well plates (ThermoFisher Scientific, Waltham, MA USA) and each plate was subsequently sterilized. PDMS coverslips were then coated with Matrigel^®^ (500 μg/mL; BD Biosciences, San Jose, CA) before plating human iPSC-derived cardiomyocytes. 50,000 (96-well format for efficacy screening) or 200,000 cells (6-well format used for drug-induced cardiotoxicity screening) were thawed, plated per well, and allowed to form functional syncytia. Cells were maintained in RPMI (ThermoFisher Scientific, Waltham, MA USA) media supplemented with B27+ (ThermoFisher Scientific, Waltham, MA USA). Cardiomyocyte functional syncytia were maintained in culture for 7 to 10 days at 37°C, in 5% CO_2_ before the screening assays. The expression of Ca^2+^ handling proteins in human iPSC-derived cardiomyocytes was determined by Western Blot analysis; a detailed description of these methods can be found in the Supporting Information.

### High-resolution optical mapping experiments in human iPSC-derived cardiac cells

We evaluated the effects of SERCA2a activators CDN1163, CP-154526, and Ro 41-0960 (Sigma-Aldrich, St Louis, MO, USA) using cryopreserved human iPSC-derived cardiomyocytes (iCell^2^ cardiomyocytes, FUJIFILM Cellular Dynamics, Madison, WI, USA). Cardiomyocyte functional syncytia were loaded with the Ca^2+^-sensitive dye Rhod-2 (ThermoFisher Scientific, Waltham, MA USA) to quantify compound effects on intracellular calcium flux, or with the voltage-sensitive dye FluVolt™ (ThermoFisher Scientific, Waltham, MA USA) to quantify changes in the membrane potential of iPSC-derived cardiac cells ^63^. Small molecules were solubilized in DMSO at a stock concentration of 10 mM. The stock concentrations of the molecules were diluted in medium to 10X the final concentration for the assay. Baseline optical mapping data was collected prior to drug treatment. Following, 96-well drug delivery plates were treated with the compounds diluted in Hank’s balanced salt solution (SAFC Biosciences, Lenexa, KS, USA) for 30 min before data acquisition. The plate was then positioned below a high-speed CCD camera (200 fps, 80 × 80 pixels) on a heating block to enable recording at physiological temperature (37 °C). Each 96-well plate was imaged using a validated LED illumination system that enables rapid acquisition of concentration-response data in a high-throughput fashion ^37^. Based on the available in situ EC_50_ data acquired in this study, we tested CDN1163 and CP-154526 for stimulation of intracellular Ca^2+^ dynamics on a 96-format well at compound concentrations of 0.1, 0.5, 1, 5, 10, and 50 µM (*n*=6 per concentration), and Ro 41-0960 at compound concentrations of 0.5, 1, 5, 10, 50, and 100 µM (*n*=6 per concentration). Additionally, we evaluated the effects of CDN1163 on conduction velocity and APD_90_ on a 6-well plate format at compound concentrations of 0.5, 5, and 50 µM (*n*=6 per concentration) to screen for potential cardiac toxicity effects induced by this molecule ^37,64^. In all cases, data acquisition occurred before (baseline) and after treatment of cells with the compounds. Post-acquisition data analysis of human iPSC-derived cardiomyocyte screening assays was done using custom software ^65,66^.

## Acknowledgments

This work was supported by the National Institutes of Health grants R01HL148068, R01GM120142 and R44ES027703 and by a Michigan Drug Discovery grant MDD2019202. This research was supported in part through computational resources and services provided by Advanced Research Computing at the University of Michigan, Ann Arbor, Michigan.

## Author Contributions

L.M.E.-F. and T.J.H. designed research; R.A.-O., J.C., E.N.J.-V., G.G.-S., N.W., and A.M. da R. performed research; R.A.-O., E.N.J.-V., G.G.-S., A.M. da R., T.J.H., and L.M.E.-F. analyzed data and interpreted results; L.M.E.-F. wrote the paper.

