## Supplementary materials for "From atoms to cells: bridging the gap between potency, efficacy, and safety of small molecules directed at a membrane protein"


**This PDF file includes:**


Supplementary Methods
Figures S1–S3
Table S1

### Supplementary Methods

**Expression of $Ca^{2+}$-handling proteins in human iPSC-derived cardiomyocytes using Western Blotting.** Cells from the 96 well plates were collected using 2x Laemmli sample buffer (Bio-Rad, Hercules, CA USA), 30 µl per well, and stored at –20°C. Samples were loaded into 4-20% Tris-Glycine polyacrylamide precast gels (ThermoFisher Scientific, Waltham, MA USA) and electrophoresis was carried out. The SDS-PAGE resolved proteins were transferred to iBlot®stacks with regular PVDF membranes using the iBlot™ 2 Dry Blotting System (ThermoFisher Scientific, Waltham, MA USA). Nonspecific binding sites were blocked with 5% nonfat dry milk in PBS-T (in mmol/L, 3 $KH_2PO_4$, 10 $Na_2HPO_4$, 150 NaCl, 0.15% Tween 20, pH 7.2-7.4) for 30 min at room temperature. Membranes were then incubated with specific primary antibodies (**Table S1**) diluted in 5% bovine serum albumin in PBS-T overnight at 4°C. After washing 3 times for 10 min, membranes were incubated with horseradish peroxidase-conjugated secondary antibodies (Table S1) diluted in 5% bovine serum albumin in PBS-T. After washing 3 times for 10 minutes, protein-antibody reactions were detected using Pierce SuperSignal Chemiluminescent Substrates (ThermoFisher Scientific, Waltham, MA USA). Detection and quantification of protein bands were performed with a Bio-Rad ChemiDoc system and Image Lab software 5 (Bio-Rad, Hercules, CA USA).

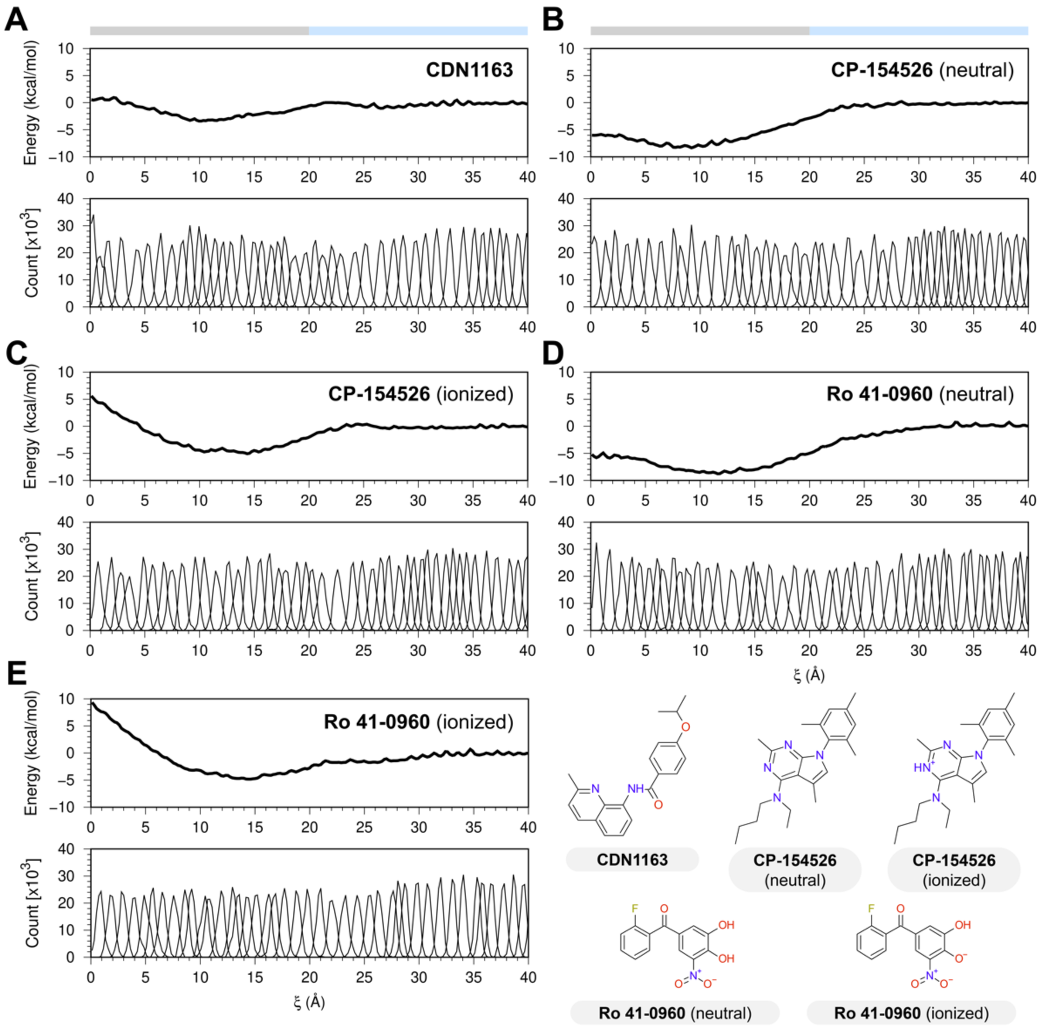


**Fig. S1. Umbrella sampling analysis.** Potential of mean force (PMF) profile (top) and histograms (bottom) of (A) CDN1163, (B, C) CP-154526, and (D, E) Ro 41-0960 through the POPC lipid bilayer along the reaction coordinate (ξ). The lower right section shows the chemical structures of the compounds and the protonation states used during the study.

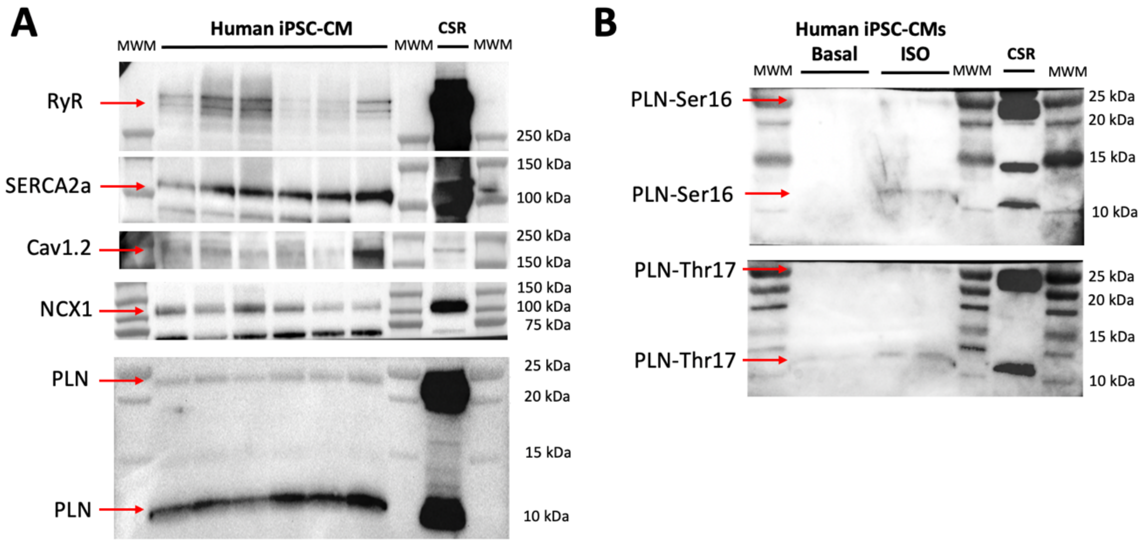


**Fig. S2. Expression of major $Ca^{2+}$ handling proteins in human iPSC-derived cardiomyocytes (iPSC-CM) by western blot analysis**. (A) Western blot analysis of iCell$^2$ cardiomyocytes showed that these cells express all major proteins involved in $Ca^{2+}$ transport and regulation of the cardiac calcium pump. We analyzed the expression of the following proteins: RyR, ryanodine receptor; SERCA2a, the cardiac isoform of sarcoplasmic reticulum $Ca^{2+}$-ATPase; Cav1.2, Voltage-dependent L-type $Ca^{2+}$ channel, α-1C subunit; NCX1, $Na^{+}$-$Ca^{2+}$ exchanger; PLN, phospholamban. (B) Detection of phosphorylated phospholamban (PLN) at positions Ser16 (PLN-Ser16) and Thr17 (PLN-Thr17) in response to pharmacological β-adrenergic stimulation with isoproterenol (ISO, 1 μM); untreated cells were included in the analysis (basal, negative control). In all cases, we used a pig heart membrane preparation as a control ('CSR' lane) for the expression of $Ca^{2+}$ handling proteins tested in this study.

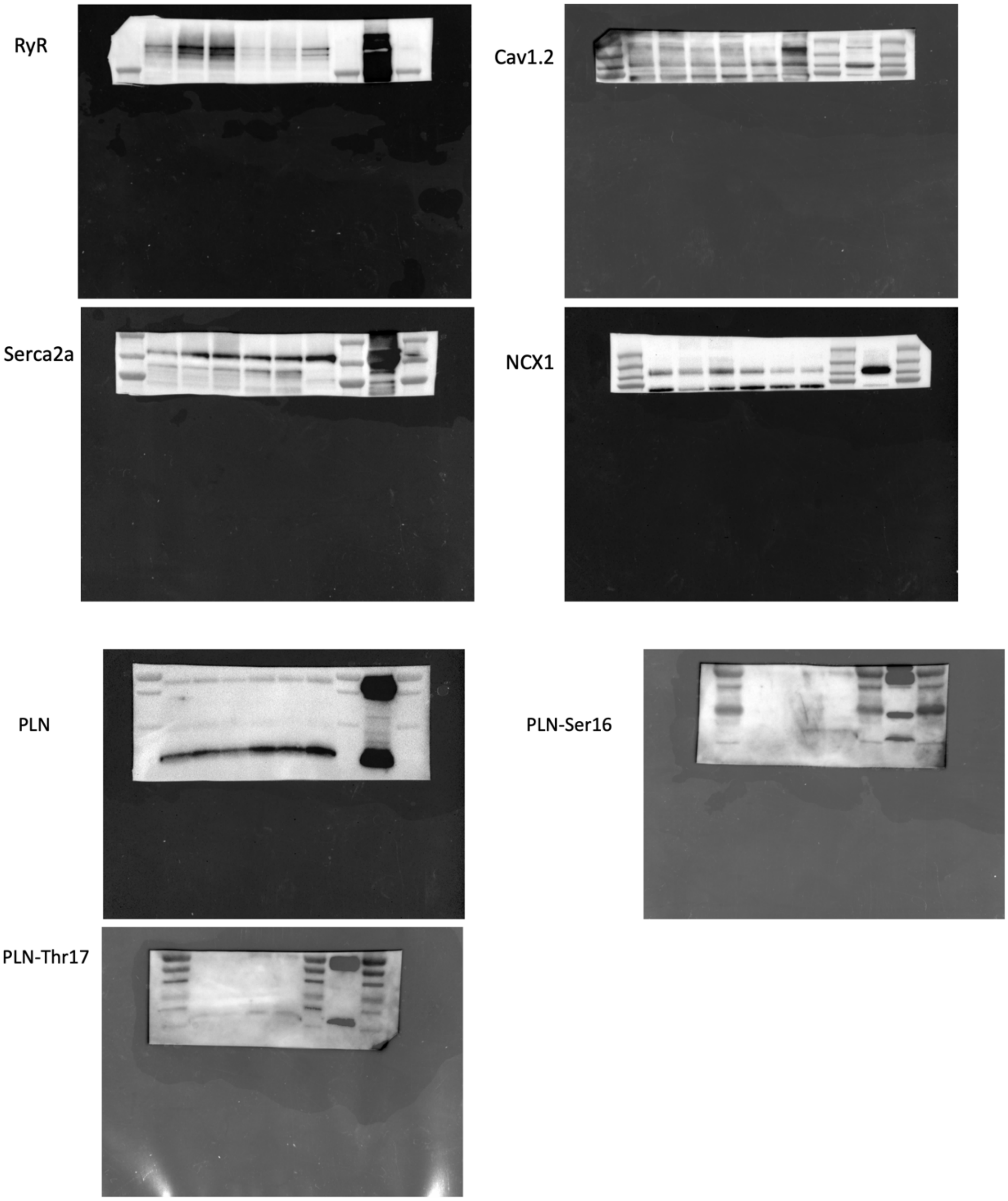


**Fig. S3.** Original images (tiff format) exported from the Image Lab software (Bio-Rad) obtained by the scanning of blots. For the final figure S2, we crop the black/grey area of each image and adjust the size in a way that all blots have the same length. In some cases, the levels were adjusted so that the bands become more apparent; the whole horizontal lines of bands were adjusted at the same time.

**Table S1.** Antibodies used to detect and quantify expression of $Ca^{2+}$-handling proteins in iPSC-derived cardiac cells

| Primary antibodies | Vendor/Supplier | Catalog # | Dilution |
|---|---|---|---|
| GAPDH | Sigma | G-8785 | 1:2000 |
| Serca2a | ThermoFisher | MA3-919 | 1:500 |
| Phospholamban | Badrilla | A010-14 | 1:5000 |
| Phospho-phospholamban (PLN-Ser16) | Badrilla | A010-12AP | 1:2500 |
| Phospho-phospholamban (PLN-Thr17) | Badrilla | A010-13AP | 1:1000 |
| Ryanodine Receptor (all RYR) | ThermoFisher | MA3-925 | 1:500 |
| Cav1.2 | Alomone | ACC-003 | 1:250 |
| NCX1 | Swant | R3F1 | 1:500 |
| **Secondary antibodies** | | | |
| Goat anti-mouse-HRP | Jackson ImmunoResearch | 115-035-166 | 1:1000<br>1:5000 for GAPDH |
| Goat anti-rabbit-HRP | Jackson ImmunoResearch | 111-035-144 | 1:1000 |
